# Spatial variation in red deer density in relation to forest disturbance and ungulate management in a transboundary forest ecosystem

**DOI:** 10.1101/2022.09.04.506509

**Authors:** Mahdieh Tourani, Frederik Franke, Marco Heurich, Maik Henrich, Tomáš Peterka, Cornelia Ebert, Julian Oeser, Hendrik Edelhoff, Cyril Milleret, Pierre Dupont, Richard Bischof, Wibke Peters

## Abstract

**Context:** Forests in Europe are exposed to increasingly frequent and severe disturbances. The resulting changes in the structure and composition of forests can have profound consequences for the wildlife inhabiting them. Moreover, wildlife populations in Europe are often subjected to differential management regimes as they regularly extend across multiple national and administrative borders. The red deer (*Cervus elaphus*) population in the Bohemian Forest Ecosystem, straddling the Czech-German border, has experienced forest disturbances, primarily caused by windfalls and bark beetle (*Ips typographus*) outbreaks during the past decades. To adapt local management strategies to the changing environmental conditions and to coordinate them across the international border, reliable estimates of red deer density and abundance are highly sought-after by policymakers, wildlife managers, and stakeholders.

**Approach and Methods:** Covering a 1081-km^2^ study area, we conducted a transnational non-invasive DNA sampling study in 2018 that yielded 1578 genotyped DNA samples from 1120 individual red deer. Using spatial capture-recapture models, we estimated total and jurisdiction-specific abundance of red deer throughout the ecosystem and quantified the role of forest disturbance and differential management strategies in shaping spatial heterogeneity in red deer density. We hypothesized that (a) forest disturbances provide favourable habitat conditions (e.g., forage and cover), and (b) contrasting red deer management regimes in different jurisdictions create a differential risk landscape, ultimately shaping density distributions.

**Results:** Overall, we estimated that 2851 red deer (95% Credible Intervals = 2609 - 3119) resided in the study area, with a relatively even overall sex ratio (1406 females, 1229 - 1612 and 1445 males, 1288 - 1626). The average red deer density was higher in Czechia (3.5 km^-2^, 1.2 - 12.3) compared to Germany (2 km^-2^, 0.2 - 11). The effect of forest disturbances on red deer density was context dependent. Forest disturbances had a positive effect on red deer density at higher elevations and a negative effect at lower elevations, which could be explained by partial migration and its drivers in this population. Density of red deer was generally higher in management units where hunting is prohibited. In addition, we found that sex ratios differed between administrative units and were more balanced in the non-intervention zones.

**Synthesis and applications:** Our results show that the effect of forest disturbances on wild ungulates is modulated by additional factors, such as elevation and ungulate management practices. Overall density patterns and sex ratios suggested strong gradients in density between administrative units. With climate change increasing the severity and frequency of forest disturbances, population-level monitoring and management are becoming increasingly important especially for wide-ranging species as both wildlife and global change transcend administrative boundaries.

## 1. Introduction

Transboundary wildlife populations are the norm in a politically divided world, rather than the exception (Liu et al 2020, Mason et al 2020). Such populations are usually subjected to spatially variable management regimes associated with separate jurisdictions. At the same time, they are often under the influence of near-ubiquitous disturbances brought on by rapid environmental changes that transcend administrative boundaries. As pressures on ecosystems mount, natural resource managers and policymakers are encouraged to seek and consider information about population-level processes (Bischof et al. 2020a). This can be exceedingly difficult to accomplish, for technical and political reasons (Bischof et al. 2016, Linnell et al 2016).

In recent decades, forest disturbances have been increasing in Europe due to alterations to forest structure and composition in combination with climatic change (Senf & Seidl 2021). Chief among them are windfalls and bark beetle (*Ips typographus*) outbreaks (Seidl et al. 2016). Bark beetles are of particular concern due to their capacity for causing extensive tree die-offs and economic damage by interrupting the transition of water and nutrients within affected trees (Seidl et al. 2008, Seidl et al. 2011, Kausrud et al. 2012). Insect outbreaks are closely related to climate change and the Norway spruce (*Picea abies*) monocultures in Central Europe (Seidl et al. 2017, Marini et al. 2017). These disturbances could impact herbivores through a change in resource availability (Pickett and White 1985) as forest openings provide enhanced foraging opportunities for many species including browsers and grazers (Kuijper et al. 2009, Ivan et al. 2018, Lehnert et al 2013, <u>Przepióra et al. 2020</u>). For example, forest disturbances due to logging and fire can increase the nutritional value of deer forage (Hayes et al. 2022).

The red deer (*Cervus elaphus*) population in the Bohemian Forest Ecosystem spans several administrative boundaries, where it experiences different levels of habitat disturbances and management interventions. While forming a contiguous population along the Czech-German border, red deer are exposed to different forms of management in the constituent national and subnational jurisdictions of the system. In the non-intervention zones of the national parks, forests are allowed to recover after disturbances without human intervention in contrast to the state forest areas and the periphery of the national parks (Zeppenfeld et al. 2015). With longstanding tradition, several winter enclosures and open feeding sites are located across the Bohemian Forest Ecosystem with regular provisioning of supplementary forage from the first snowfalls until green-up in spring (Rivrud et al. 2016a). In spring, forage provisioning is stopped at open feeding sites, enclosures are opened, and deer can move freely during the growing season. This management tool is used to encourage deer to stay in designated wintering sites to prevent bark stripping and browsing, to help regenerate forest (Möst et al. 2015). Deer are counted every winter in the different enclosures and these annual counts have been traditionally used as an index for changes in relative abundance. However, with ongoing climatic change and thus milder winters, more deer may spend winters outside the fenced enclosures and away from open feeding sites. Policymakers, local managers, and other stakeholders involved in red deer management in the Bohemian Forest Ecosystem have been seeking information about the status of the population, especially estimates of abundance, and learning about the role that forest disturbance may play in red deer density and distribution.

Here, we used non-invasive faecal DNA sampling and spatial capture-recapture (SCR) analysis to (a) estimate density distribution of red deer throughout the Bohemian Forest Ecosystem, and (b) test for the effects of forest disturbances and management regimes on red deer density throughout this transboundary ecosystem. The forage maturation hypothesis proposes that ungulate migration is driven by selection for high forage quality (Rivrud et al. 2016b). Therefore, we expected red deer densities in summer to be higher in areas with greater forest disturbances, primarily because of higher food availability (Kuijper et al. 2009) due to an abundance of early seral stands. We also expected the positive effect of forest disturbances on red density to be stronger at higher elevation as human interventions on forest disturbances are generally lower in these areas. We expected local management practices to affect red deer density distribution and that the highest abundance occurs in protected areas where hunting pressure is the lowest.

## 2. Methods

### 2.1. Study area

Our study area straddled the Czech-German border, covering three different administrative units in the Bohemian Forest Ecosystem: (1) the Bavarian Forest National Park (BFNP, 245 km^2^) and (2) the State Forest Neureichenau (SFNR, 152 km^2^) in Germany, and the majority of (3) the Šumava National Park (SNP, 684 km^2^) in Czechia (Fig. 1). The two parks are characterized by intermediate elevations with several mountains along the border between Germany and Czechia. The parks are surrounded by low-elevation managed forests, such as SFNR in Germany and the military training area Boletice and the state forest district Boubin on the Czech side (Fig. 1), which are part of regions with lower protection status on both sides of the border (BFNP and the Bohemian Forest Protected Landscape Area; Heurich et al. 2015). Both protected landscapes, neighbouring with national parks, form natural buffer zones. The red deer habitat is restricted by law on the German side to an approximately 604-km^2^ area that is only marginally larger than BFNP and SFNR. Outside of this designated red deer area, all red deer should be culled by law during the regular hunting season. In Czechia, however, the red deer occurrence does not have such a solid border and continues further into the neighbouring Bohemian Forest Protected Landscape Area. The non-intervention zone of the national parks prohibits hunting (herein, no-hunting zone).

**Figure 1.**
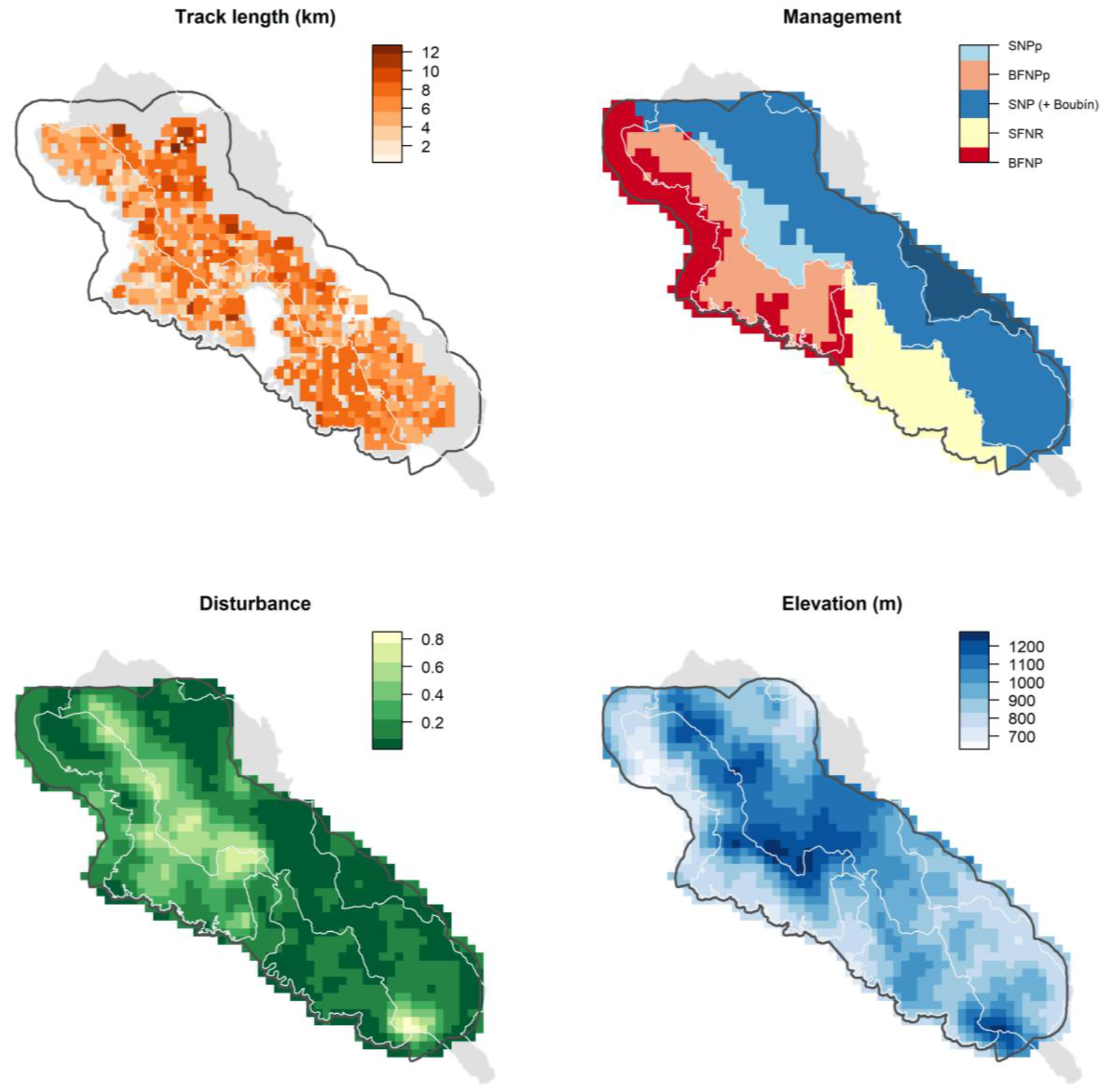
Spatial depiction of covariates included to model variation in red deer detection probability (top-left panel: recorded GPS search-tracks) and density (from top-right panel, clockwise: a. management units, b. elevation, c. proportion of forest disturbances). The management units are: (1) BFNPp: the no-hunting zone of the Bavarian Forest National Park, (2) BFNP: ungulate management zone of the Bavarian Forest National Park, (3) SFNR: the State Forest Neureichenau district, (4) SNPp: the no-hunting zone of Šumava National Park, and (5) SNP: the ungulate management zone of the Šumava National Park and a small part of the State Forest district Boubín in the buffer of the central part of the study area on the Czech side (darker blue area outside the white lines). For visibility, soft edges of the management units are not shown. White and black lines show the three administrative units and the border of the study area used in the spatial capture-recapture analysis, respectively.

With an elevation range from 570 to 1453 m, the study area is dominated by coniferous forests (60%), mixed forests (20%), and grasslands including pastures (14%). Several open areas within the forest were created by bark beetle outbreaks (Pflugmacher et al. 2019), which first occurred in 1983, reached a peak in 1996-1997 (Lausch et al. 2011), and have continued since. Other land cover types present in the area include broadleaved forests (6%), shrublands (< 1%), and surfaces covered with buildings and extensive pavement (< 1%; Pflugmacher et al. 2019). Forests in the study area are mainly comprised of Norway spruce in moist valleys and transition to mixed forests with an abundance of European beech (*Fagus sylvatica*) and silver fir (*Abies alba*) at intermediate elevations. Forests at high elevation are less dense and rich in Norway spruce, interspersed with mountain ash (*Sorbus aucuparia*) and sycamore (*Acer pseudoplatanus*). In SNP, Norway spruce has replaced natural forests in more extensive areas than in the BFNP (Krojerová-Prokešová et al. 2010, Cailleret et al. 2014).

Red deer dominates the ungulate guild in the study area, but roe deer (*Capreolus capreolus*) also occurs, albeit at lower densities. Wild boar (*Sus scrofa*) is more common on the Czech than on the German side, but distributions fluctuate. Fallow deer (*Dama dama*) and moose (*Alces alces*) seldom traverse the study area (Janik et al. 2021). The western part of Czechia is also inhabited by the non-native sika deer (*C. nippon*) that are regularly culled in the study area (Saggiomo et al. 2021). However, the threat of hybridization between sika and red deer remains unmanaged (Bartoš 1981, Krojerová-Prokešová et al. 2017). Red deer in the study area are partially migratory (Peters et al. 2019). Natural predators include wolves (*Canis lupus*), with a first pair having recolonised the area in 2016, and Eurasian lynx (*Lynx lynx*), which occasionally prey on red deer females and calves (Belotti et al. 2014). Humans pose the main predation risk for red deer in the Bohemian Forest Ecosystem. Hunting quota are set based on annual counts in winter enclosures and at feeding stations, number of red deer hunted in the previous year, and an inventory of browsing damage (Möst et al. 2015). Red deer are hunted mainly from high stands by single hunters or small groups of hunters in autumn, while in late autumn and winter drive hunts outside BFNP may also be practiced. In SFNR, red deer are hunted under regular German and Bavarian hunting guidelines. In SNP, hunting is prohibited in approximately 10% of the area. Hunting is prohibited in 75% of BFNP and a large proportion of red deer culling occurs when the animals enter the winter enclosures. The hunting season lasts from June to January in Germany and from August until mid-January (until end of March for calves) in Czechia. Overall, as of 2018, hunting was prohibited in 23% of the study area (Fig. 1), and in 19% of the study area hiking was restricted to marked trails to reduce human disturbances.

### 2.2. Faecal DNA sampling and genotyping

To guide non-invasive DNA sampling of the red deer population, a 1-km^2^ grid was generated using ArcGIS Desktop 10.5.1. Due to the large extent of the study area, not all grid cells could be searched and following a simulation study of sampling design trade-offs, we randomly discarded 20% of the grid cells, but avoided discarding neighbouring grid cells to limit the size of resulting spatial gaps in sampling. We also omitted 42 grid cells because more than half of their area was covered by human settlements, water bodies, or very steep terrain that was difficult to access by searchers. The final search area included 543 grid cells (Fig. 1).

We collected fresh deer faeces between 1 June and 26 July 2018 (49 sampling days). Surveyors conducted structured search-encounter sampling and GPS-recorded their search-tracks and location of samples. To ensure homogeneous coverage, each grid cell was subdivided into 16 smaller units of 250 × 250 m, which were searched with similar intensity (Fig. 1). Surveyors were advised to walk about 10 km within each 1 km^2^ grid cell. However, steep terrain that was dangerous to access was skipped. We collected only fresh pellets, but in areas with particularly low numbers of detections, older pellets with a relatively intact surface were also sampled. We further enforced a minimum distance of 2 m between subsequent pellet groups to avoid sampling the same group twice. From each pellet group, two individual pellets were sampled. A new toothpick or latex glove was used for every sample to avoid cross-contamination when transferring the sample into a falcon tube. At the end of each day, samples were placed in a freezer at −20°C.

Genetic analyses were performed according to Ebert et al. (2021). Briefly, after DNA extraction, eight dinucleotide microsatellites and one sex marker (Gurgul et al. 2010) were amplified in two multiplex PCRs (Table S1). Two negative controls were included in all PCRs to detect potential contamination. Determination of matching genotypes was carried out using GENECAP (Wilberg and Dreher 2004). We scrutinized genotypes differing by one (1-MM) or two (2-MM) alleles to detect genotyping errors. For all 1-MM and 2-MM pairs, raw data were re-checked to resolve the mismatches. Genotype pairs with only one mismatch were regarded as originating from the same individual (Ruell et al. 2009). Those pairs with 2-MM were considered as originating from different individuals if re-checking of raw data and an additional two PCR repeats did not alter the results and if both samples matched with other samples in the data set (Paetkau 2003). To confirm the power of the used loci, we calculated the probability of identity and the probability of identity for siblings as a more conservative metric (Waits et al. 2001) and heterozygosity using GIMLET (Valière 2002). We calculated genotyping error rates (allelic dropout and false alleles) as recommended in Broquet & Petit (2004; equations 1 and 3). Samples identified by their genotype as originating from roe or fallow deer were excluded from further analyses. As a result, the data consisted of individual identity, sex, and location associated with non-invasive red deer detections.

### 2.3. Analysis

We built an SCR model in a Bayesian framework (Royle 2009) with two hierarchical levels distinguishing the observation process from the ecological process (see Supporting Information for model definition).

#### i) Ecological process

In SCR models, individual locations are defined by their centre of activity or home range. Abundance is then defined as the number of individual activity centres within the region of interest or habitat. Here, we defined the habitat as the area searched for red deer DNA samples surrounded by a 3-km buffer to account for edge effects (Efford 2011) leading to a habitat polygon of 1209 km^2^ subdivided in 1 *×* 1 km grid cells (Fig. 1). In SCR analysis, the buffer allows to explicitly account for the possibility to detect individuals that had their activity centre outside of the searched area (Efford 2004). To explore the drivers of red deer density, we modelled the distribution of activity centres as an inhomogeneous Bernoulli point process (Illian 2008, Zhang et al. 2022) whose intensity is proportional to the red deer density and related to a set of covariates:

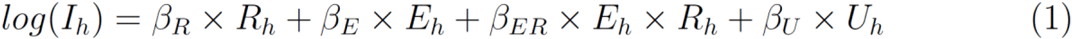

In this formulation, *I*_*h*_ is the point process intensity in habitat grid cell *h*, and *R*_*h*_, *E*_*h*_, and *U*_*h*_ are covariates describing the proportion of disturbed habitat, the average elevation, and management unit within a 2-km radius of habitat grid cell *h*, respectively. *β* coefficients are the effects of covariates (see below) in a given habitat cell on the probability that an individual has its activity centre located in this same cell. The main effects include the proportion of forest disturbances *β*_R_, elevation *β*_E_, and management region *β*_U_ (Eq. 1). As disturbances tend to occur at different elevations (e.g., windthrows predominantly at higher elevations; Oeser et al. 2017), and elevation as proxy for plant phenology has been shown to be a main predictor for red deer migration (Rivrud et al. 2016a) and habitat selection (Heurich et al. 2015), we included an interaction between elevation and forest disturbances. *β*_ER_ is the interaction terms of disturbance and elevation. In addition, because management regimes differ amongst and within the different administrative units, we considered that red deer density could differ between the five management regions, corresponding to three administrative units and their hunting/no-hunting zones (Fig. 1). *β*_*U*_ is the slope for the management region of the habitat cell and is compared to the ungulate management zone of SNP (intercept), where hunting is authorized (Fig. 1).

Proportion of disturbance was generated using the Landsat-based forest disturbance maps created by Oeser et al. (2017). We combined the categories provided by Oeser et al. (2017), i.e., bark beetle infestations, windthrows, salvage-logged wind throws and bark beetle sites. We derived elevation from Shuttle Radar Topographic Mission (SRTM) maps downloaded at 1 × 1-km resolution. Land protection status was provided by the national parks. Management units include: (1) ungulate management zone of BFNP, (2) non-intervention zone of BFNP, (3) ungulate management zone of SNP, (4) non-hunting zone of SNP, and (5) SFNR (Fig. 1). We considered a moving window around each habitat cell size to calculate the proportion of each management unit within a 2 km radius, where the values gradually decreased towards the edges from 1 to 0. All spatial covariates were resampled to the habitat resolution (Fig. 1), then standardised before model fitting.

To account for the fact that some individuals in the population may never be detected, we used a data-augmentation approach (Royle et al. 2007). Following this approach, we derived estimates of population size *N* by summing the number of individuals included in the population, where *M* is the maximum possible number of individuals in the population.

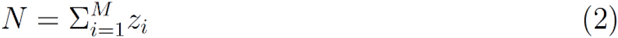

We modelled individual inclusion in the population through a latent state variable *z*_i_, governed by the inclusion parameter Ψ for all individuals *i* in 1:*M* as:

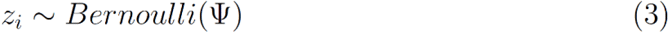

#### ii) Observation process

The SCR observation component models how the individual detection probability varies over a set of detectors. Here, we generated detector locations by discretizing the search area into 4122 grid cells of size 400 *×* 400 m (Fig. 1). We used the partially aggregated binomial model to retain as much information from the collected genetic data as possible (Milleret et al. 2018) and further divided each detector grid cell into 16 sub-cells (or less if some sub-cells did not overlap the suitable red deer habitat based on our knowledge of the study system). We then generated individual spatial detection histories by retrieving the frequency of sub-cells with at least one sample from the focal individual for each detector main grid cell.

We used the half-normal detection function (Royle et al. 2014) and modelled the probability of detecting an individual at a given detector as a decreasing function of the distance between this individual’s activity centre and the detector:

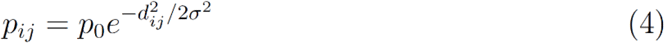

Here, *p*_*ij*_ is the detection probability of individual *i* at detector *j, p*_0_ is the baseline detection probability, *d*_*ij*_ is the distance between individual *i*’s activity centre and detector *j*, and *σ* is the scale parameter which dictates how fast the detection probability decreases with distance. To account for spatial variation in detectability, we modelled a detector-specific baseline detection probability:

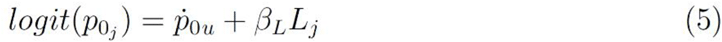

In this equation, *p*_0*u*_ is a separate baseline detection probability for each of the three administrative units (BFNP, SFNR, and SNP) because of the potential variation in sampling effort. *L*_*j*_ is the length of GPS search tracks recorded within detector grid cell *j* and *β*_*L*_ is the slope parameter describing the linear relationship between effort and detection probability.

#### iii) Model fitting and post-processing

We fitted sex-specific models using NIMBLE version 0.6-9 (de Valpine et al. 2017) and R (version 3.5.2, R Development Core Team 2018) with functions now available in the R package nimbleSCR (Bischof et al. 2020b). We ran 4 chains of 100,000 iterations each and discarded the first 10,000 samples as burn-in, leading to a total of 360,000 MCMC samples per model to draw inferences from. We assessed convergence by looking at the potential scale reduction value for all parameters and mixing of the chains using trace-plots (Brooks and Gelman 1998). For mapping density, we thinned the posterior samples by 10 and based the maps on 36,000 samples. To obtain estimates of abundance for each administrative unit, we summed the number of model-predicted activity centres that fell within the administrative unit of interest for each iteration of the MCMC chains, thus generating a posterior distribution of the abundance for this area, from which mean abundance estimates can be derived. For prediction of density as a function of covariate effects, we calculated the relative density per cell and multiplied the habitat intensity value by the *N* estimate in each cell for every MCMC iteration.

## 3. Results

### 3.1. Faecal DNA sampling and genotyping

During sampling, 3450 km of GPS search tracks were recorded, and 3234 putative red deer faeces were collected. The 1578 (48.8%) successfully genotyped samples were assigned to 1120 red deer individuals (494 females, 560 males, and 66 of unknown sex due to amplification failure of the sex marker). Of the genetically identified individuals that were included in the analysis (n = 1054), 28.5% were detected more than once (33.7% of the detected males and 25.9% of the detected females) with a maximum of six samples from the same individual. Genotyping error rates are reported in Table S2. The mean allelic dropout rate over all loci was 4.2%, whereas the mean false alleles rate was 0.9%. Overall Probability of Identity of the data set was 1.89 × 10^−11^, and the overall probability of identity for siblings was 0.00016.

### 3.2. Abundance and density estimates

We estimated red deer population size in our study area at 2851 individuals (95% Credible intervals CI = 2609 to 3119). Sex-specific estimates were 1406 females (95% CI = 1229 to 1612) and 1445 males (95% CI = 1288 to 1626) for the summer of 2018 (Fig. 2, Table S3). The overall sex ratio was even (F:M = 1:1.03), but differed between management zones, with a slight skew towards males in SNP (1:1.07), a strong skew towards males in the SFNR (1:2.06) and a female bias in the BFNP (1:0.78). The overall abundance was higher on the Czech side (2052 red deer, 95% CI = 1836 to 2292) compared to the German side (800 red deer, 95% CI = 680 to 940). Likewise, the average red deer density was higher in Czechia (3.5 km^-2^, 1.2 to 12.3) compared to Germany (2 km^-2^, 0.2 to 11; Fig. 2).

**Figure 2.**
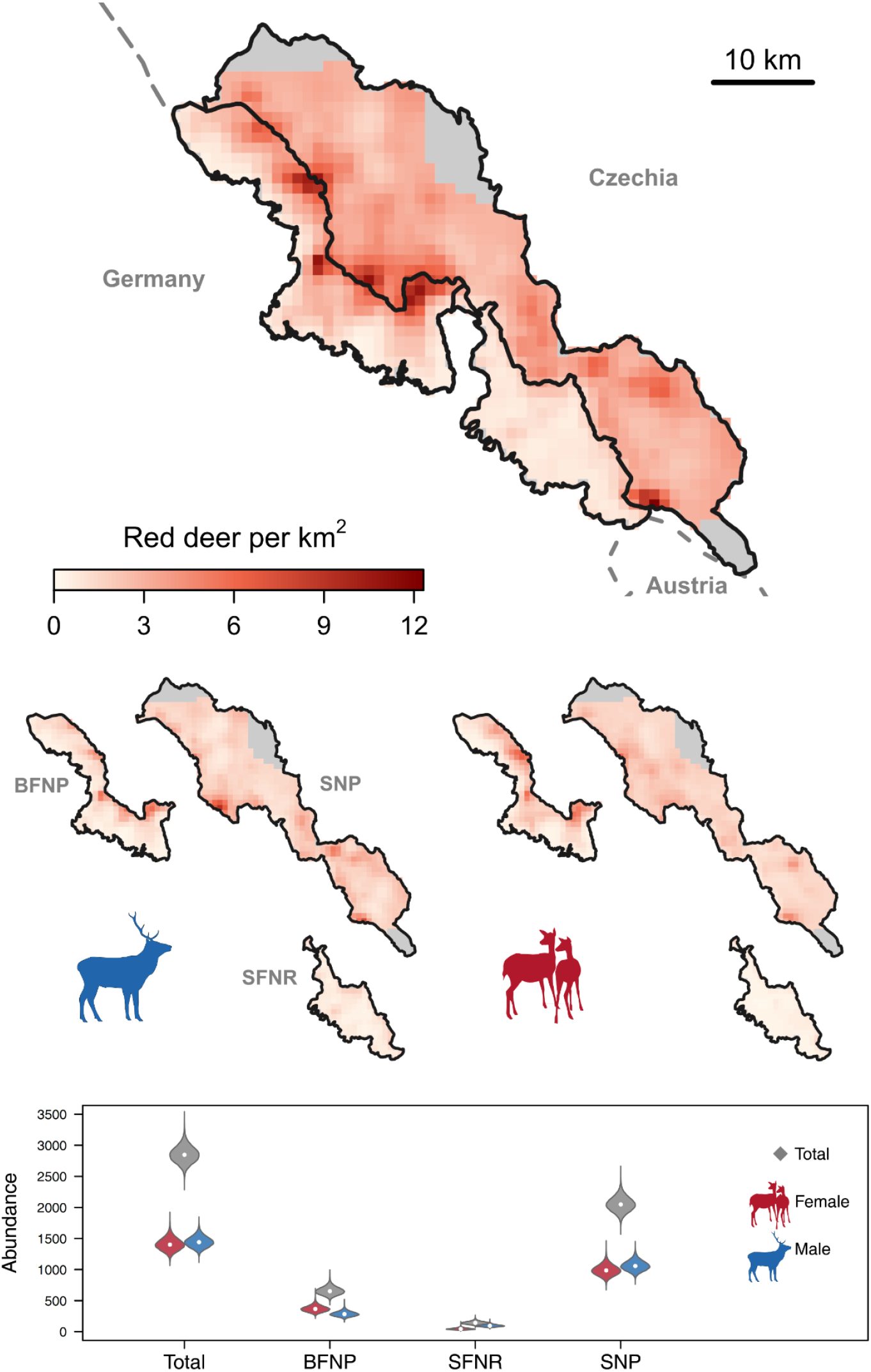
Realized density maps (upper panels) and abundance estimates (lower plot) for red deer *Cervus elaphus* across the Bohemian Forest Ecosystem in summer, June, and July 2018. Population estimates are broken down into sex-specific estimates for the three administrative units (BFNP: Bavarian Forest National Park, SFNR: State Forest Neureichenau, and SNP: Šumava National Park). Gray areas in density maps represent regions beyond the sampled extent that belong to the management jurisdictions. Violins in the lower plot show posterior distributions of abundance with 95% credible interval and white dot indicates the medians.

We estimated the average red deer density for the entire study area in the Bohemian Forest Ecosystem at 1.42 females and 1.46 males km^*−*2^. The effect of forest disturbances on red deer density was modulated by elevation, changing from negative at low elevations to strongly positive at high elevations (Fig. 3; Table S4). The ungulate management zone of BFNP and SFNR on the German side had lower baseline red deer densities compared to the ungulate management zone of SNP on the Czech side (Female *β*_BFNP_ = −2.8, 95% CI = −4.8 to −1.4 and Male *β*_BFNP_ = −2.2, 95% CI = −3.8 to −1; Female *β*_SFNR_ = −1.9, 95% CI = −2.7 to −1.1 and Male *β*_SFNR_ = −1, 95% CI = −1.5 to −0.5; Table S4). The non-intervention zone of BFNP had a higher baseline density, compared to ungulate management zone of SNP as the reference area, but beta coefficients overlapped zero (Table S4). The non-intervention zone of SNP had lower red deer densities compared to the management zone of the SNP, but coefficients overlapped zero (Table S4).

**Figure 3.**
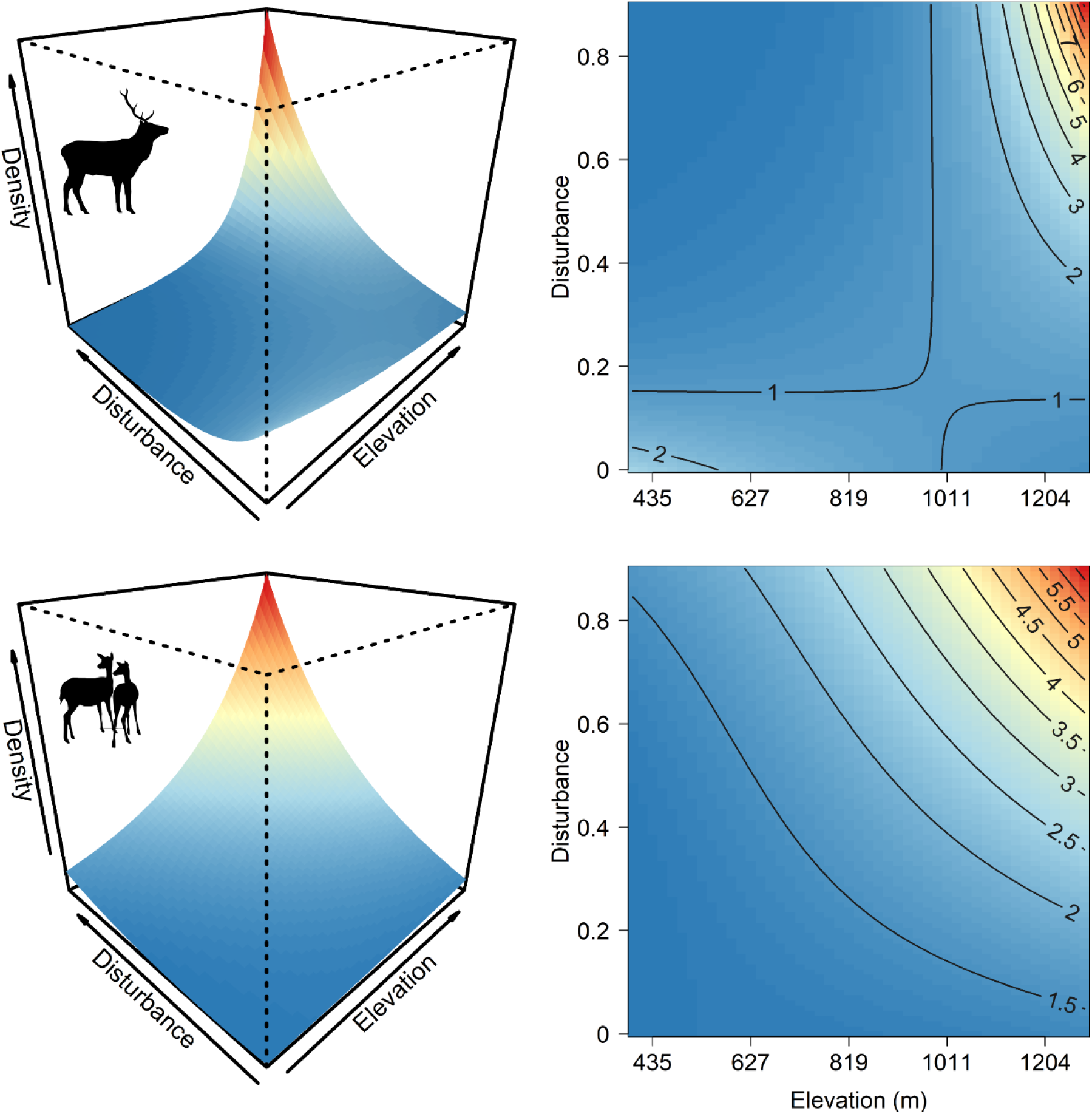
Sex-specific predictions of red deer relative density (individual per km^-2^) across the Bohemian Forest Ecosystem as a function of the interaction between proportion of forest disturbances and elevation (top: male, bottom: female deer). Contour lines in the plots to the right represent relative density of red deer.

Detection probability was positively associated with the length of transects searched for both sexes (Female *β*_*L*_ = 0.4, 95% CI = 0.3 to 0.5 and Male β_L_ = 0.5, 95% CI = 0.4 to 0.6). The scale parameter of the half-normal detection function *σ* was similar between female (*σ* = 1 km, 95% CI = 0.9 to 1.1) and male red deer (*σ* = 0.9 km, 95% CI = 0.8 to 0.9). These *σ* values translate to average summer home-range sizes of 18 km^2^ (95% CI = 15 to 21) and 15 km^2^ (95% CI = 13 to 17) for females and males, respectively, during the sampling period.

## 4. Discussion

The transboundary population of red deer in the Bohemian Forest Ecosystem along the Czech-German border is characterized by higher densities at higher elevations and in areas with forest disturbances, which are associated with higher forage quality in summer. Red deer density was up to six times higher in areas subjected to forest disturbance than in undisturbed areas, especially at higher elevations (Fig. 3). Red deer density was also higher in the non-intervention zones of the protected areas, compared with areas where hunting was authorized. We also found pronounced sex-specific spatial variation in density (Fig. 2). Both forest disturbances and different ungulate management regimes across the administrative units explained this variation.

### 4.1. Context-dependent effect of forest disturbance management

While forage availability is likely driving habitat quality in forest disturbance gaps, ungulates have to trade-off between forage and predation risk (Dupke et al. 2017, Rettie and Messier 2000). The positive effect of forest disturbances on red deer density in the Bohemian Forest Ecosystem was context dependent and most pronounced for males. Specifically, while red deer seemed to avoid disturbed areas at low elevations, disturbances were attractive for red deer at high elevations (Fig. 3). Forest openings created by disturbances have the potential to provide ungulates access to diverse and abundant forage that follows the removal of overstory canopy (Kuijper et al. 2009). In addition, disturbed areas – if they stay unmanaged – provide excellent shelter. While disturbances caused by bark beetle infestations and windthrows remain unmanaged in the non-intervention zones of the national parks, which cover large parts of the high-elevation areas in the study system, disturbances in the management units of the national parks and in SFNR are managed through salvage logging. For forest disturbances, we could not distinguish windthrows and bark beetle infestations from salvage logged areas as two different treatment groups. However, salvage logged areas were a small fraction of the managed forest disturbances in the study area.

Forests recover faster and more homogeneously at salvage-logged than at non-intervention sites in the Bohemian Forest Ecosystem (Senf et al. 2019). In addition, forest recovery is faster in low elevations, leading to a long-lasting increase of habitat quality in the higher compared to the lower elevations (Lone et al. 2015, Oeser et al. 2021). Additionally, the type of vegetation covering disturbed sites differs between high and low elevations, with more grasses and ferns at high elevations, which might also be more attractive to mixed feeders like red deer (Ewald et al. 2014). Finally, at low elevations, more open habitats associated with higher hunting pressure at disturbed sites may offset the positive effect of increased forage availability, leading to red deer avoiding disturbed areas at lower elevations. This is in accordance with the pattern of increased positive association between disturbances and red deer density with elevation (Table S4). We detected similar patterns of positive elevation-disturbance effects on red deer density, but they were more pronounced for male than for female red deer.

### 4.2. Seasonal migration and forage availability

Red deer inhabiting the Bohemian Forest Ecosystem are partially migratory, i.e., only part of the population migrates, while the remainder stays resident on the shared winter range (Peters et al. 2019). Migration behaviour, and hence habitat selection, is strongly affected by forage phenology as suggested by the forage maturation hypothesis (Rivrud et al. 2016b). Specifically, individuals migrating to higher elevations in spring have access to more high-quality forage during the growing season compared to red deer that remain at lower elevations (Bischof et al. 2012). In this context, forest disturbances play a crucial role in red deer habitat selection and distribution patterns (Oeser et al. 2021), which was supported by our findings. Gaps provided by forest disturbances often increase foraging opportunities due to a higher abundance of plant biomass on the ground (Kuijper et al. 2009 but see Ewald et al. 2014). Recent studies suggest that habitat suitability for red deer improved after disturbance for at least 25 years, and these disturbance-related habitat effects generally increase with elevation (Oeser et al. 2021). Specifically, different disturbance types occurred along elevational gradients and wind throws were characteristic for higher elevations (Oeser et al. 2021). Post-disturbance recovery is also affected by elevational gradients in the Bohemian Forest Ecosystem, further affecting post-disturbance recovery (Senf et al. 2019), with salvage logging mainly occurring at lower elevations. Overall, we observed a pattern that is typical for a partially migratory ungulate population under the predictions of the forage maturation hypothesis (McNaughton 1985, Fryxell 1991, Rivrud et al. 2016b). These findings are mediated by ungulate management in our study system.

### 4.3. Management interventions and the effect of hunting

Hunting is the main selective force for the red deer in our study area, as for many transboundary ungulate populations in temperate climates (Frid and Dill 2002). Hunting has been shown to affect distribution and, hence density, of *Cervus* spp. (Ciuti et al. 2012, Rivrud et al. 2016b, Meisingset et al. 2018). In the Bohemian Forest Ecosystem, summer red deer density was the lowest in SFNR, where population size is regulated by hunting and elevation is lower compared to the two national parks (Fig. 2; Table S3). In contrast, density was more than twice as high in BFNP, which includes a large non-intervention zone. The highest densities were predicted for SNP, where deer are also protected year-round in the non-intervention zone (i.e., hunting is not authorized), but more deer were hunted outside the non-intervention zone compared to the BFNP or SFNR.

We detected a difference in red deer sex-ratio amongst the three administrative units with a 1:2 female to male ratio in SFNR, compared to 1:1.1 and 1:0.8 in SNP and BFNP, respectively. In contrast, the winter enclosure counts in SFNR rather suggest a female-biased sex ratio. During the winter preceding this study, 419 red deer with a sex ratio of 1.16:1 (F:M) in BFNP, 222 red deer (1.98:1) in SFNR, and 562 red deer (1.08:1) in SNP were counted in the enclosures or at open feeding sites (n = 1203). The current hunting regulations protects males older than three years in BFNP, but this has not resulted in a male-biased sex ratio in this area based on our findings (Table S3). The male-biased sex-ratio observed in SFNR is most likely the result of a combination of hunting focused on females to regulate population size, differential space use between males and females during summer, and a high proportion of female migrants that use SFNR in winter only. For example, there are extensive forest disturbance areas in SFNR that provide high-quality forage, yet the deer do not use this area and move to the Czech side instead, where hunting pressure is lower. Likewise, telemetry data show pronounced seasonal dynamics in space use of red deer in the Bohemian Forest Ecosystem, with a considerable proportion of female red deer that spent the winter in the enclosures in SFNR, migrate in spring eastward into the border region between SFNR and SNP, or north to the high-elevation disturbed areas between the two national parks (Peters et al., unpublished data). Migration to the open German-Czech border zone provides access to high-quality forage similar to the forage availability at disturbed areas and seems to support ideal conditions for females raising offspring. Females with calf might also prioritize risk avoidance more than males (Lone et al. 2015).

While our study produced actionable information for the transboundary red deer population, managers and decision-makers should be aware that we only report a snapshot representation of the population for a limited time frame. For example, spatial patterns in density during the hunting season, which mainly occurs in autumn, can be expected to differ from those presented here (Rivrud et al. 2016b). Most importantly, our summer abundance estimates differ from the winter counts, which can be due to seasonal movements, raising the question whether winter counts are appropriate to derive hunting quota in our study system. Our sampling period was constrained by deer migration to the summer habitats in spring and the rutting season and the migration in autumn. Alternatively, sampling that is completed after the rut, but before migration, would better represent the autumn population and distribution, which is most relevant to harvest management. Sampling in this period would also probably yield better DNA quality due to lower ambient temperatures resulting in higher genotyping success rates and would be more feasible from a practical point of view. Furthermore, the Bohemian red deer population extends over a larger area beyond the one we sampled here, especially on the Czech side, and seasonal movements into or out of our study area are likely. Integration of population-level monitoring into an adaptive management framework will require periodic monitoring and an informed choice of the seasons to be prioritized for sample collection. The limitation would be the difficulty of conducting fieldwork and cost of sample collection and DNA analysis.

### 4.4. Abundance estimates and jurisdictions

Each jurisdiction in the Bohemian Forest Ecosystem manages its part of the red deer population largely autonomously. In our study, the effect of differential management systems became visible at the border between SFNR with low red deer densities and SNP with high densities. However, when different management systems pursue a common goal, such as the two national parks in their neighbouring non-intervention zone of BFNP and the no-hunting zone of SNP, red deer densities are comparable. Our results confirmed that there is a continuous exchange of red deer between Germany and Czechia, especially in the high-elevation core zones of the two national parks. Thus, red deer on both sides of the national and subnational borders are part of one contiguous population. Coordinated population-level monitoring and analysis yielded a model-estimated distribution of activity centres (Fig. 2) – an individual-based representation of the red deer population from which abundance estimates can be extracted for any desired spatial extent and therefore at multiple spatial scales or administrative levels (Bischof et al. 2020a). The spatially explicit nature of the analysis also accounts for the fact that borders are permeable and that individuals living near them may cross them.

### 4.5. Implications beyond the study system

In multiple ways, the case of the Bohemian Forest red deer population is representative of managed transboundary wildlife populations in Europe and elsewhere. Populations of large mammals are frequently shared by multiple jurisdictions, including nations. Regardless of how much they differ in their goals and actions, management on different sides of a border becomes intertwined through its impacts on the shared population. This is not limited to ungulates; similar situations are faced by large carnivore managers in northern and Central Europe, where carnivore populations are shared by several countries (Bischof et al. 2016, Gervasi et al. 2019). Despite differences in management objectives and strategies in Germany and Czechia, we recommend coordinated monitoring and joint analysis to produce population-level estimates of abundance and density.

Population size and density are perhaps the most fundamental measures used in wildlife monitoring and for setting management goals. Yet, they can be challenging to obtain, and decision-makers often end up relying on proxies or indices of questionable reliability (Moqanaki et al. 2018, Gopalaswamy et al. 2022). Technical advancements and the rising popularity of non-invasive monitoring methods have made population-level monitoring more accessible (Tourani 2022). The resulting data, in combination with analytical methods that account for imperfect and variable detectability, can yield absolute estimates of abundance, and thus can be used to show the effects of management practices or the change of ecological processes, such as seasonal migrations or imbalanced sex ratios over time. Such methods will become increasingly important since wildlife managers are not only challenged by the administrative separation of the population, but also by the ubiquitous pressures associated with ongoing human-caused global change. Management tools introduced decades ago and successfully used in the past may lose efficiency in the future. For example, in our study landscape, the enclosure system used to manage the red deer population in winter as a damage mitigation tool might lose its efficiency by milder winters because of climate change. Likewise, the expansion of disturbed areas caused by windthrows and bark beetle outbreaks may offer emerging forage areas for red deer in winter. Thus, traditional wildlife management interventions need to be updated by evidence-based sustainable practices.

## Acknowledgments

This project was funded by the program Ziel ETZ Free State of Bavaria – Czechia 2014-2020 (INTERREG V). M.T. was supported by a postdoctoral research fellowship from Conservation International through UC Davis during the writing of this manuscript. R.B., C.M., and P.D. were partially funded by the Research Council of Norway (NFR 286886, project WildMap). We thank the field staff and volunteers that helped with collecting the faecal samples.

## Supporting Information

Additional supporting information will be made available upon acceptance.

